# Morphometric Latent Factors in Autism and Their Association with Receptor Profiles and Behavior

**DOI:** 10.64898/2025.12.08.692851

**Authors:** Anas Al-Naji, Amirhussein Abdolalizadeh, Daniel Kristanto

## Abstract

The profound heterogeneity of Autism Spectrum Disorder (ASD) is a major barrier to developing targeted therapies. While dimensional subtyping using functional connectivity (FC) has advanced the field, the intrinsic instability of FC limits its power to identify stable trait-like biomarkers. This study provides a direct comparison of latent factors derived from both FC and a stable morphometric measure, Morphometric Inverse Divergence (MIND), within the same ASD cohort. We hypothesized that stable structural factors would provide a more behaviorally relevant and biologically grounded account of ASD heterogeneity.

Our findings reveal an important dissociation between functional and morphometric features. Latent factors derived from morphometric similarity (MIND) significantly correlated with core ASD behavioral traits (SCQ, SRS), while functional factors showed no association. This dissociation was also found at the neural level. Specifically, we observed weaker structure-function correspondence in the ASD group compared to the healthy control group (HC). Moreover, the stable structural factors were linked to trait-like neurodevelopmental mechanisms: the association with the CB1 receptor (important for synaptic pruning) observed in the HC was notably missing in the ASD group. Conversely, the functional factors were associated with a state-like arousal system (the norepinephrine transporter, NET).

Collectively, our results demonstrate that stable morphometric-based factors, rather than time-varying functional ones, are predictive of behavioral traits in ASD. This work validates MIND as a robust approach and suggests that the link between structural organization and its neurodevelopmental (CB1) underpinnings is a more powerful and stable target for developing biomarkers in autism.

## 1. Introduction

Autism Spectrum Disorder (ASD) is a neurodevelopmental syndrome encompassing a range of conditions, such as difficulties in communication and social interaction, alongside repetitive and restricted behaviors (Lord et al., 2018). These conditions can create a significant burden for the diagnosed individuals and for the society that usually provides substantial caring efforts (Buescher et al., 2014). To this end, extensive research has been conducted to discover effective medications and treatments for ASD. However, although the common conditions of ASD are known, the profound heterogeneity across individuals in terms of their clinical presentation and behavioral profiles necessitates more personalized treatment and medication (Lombardo et al., 2019). Efforts to address this heterogeneity include subtyping. To date, various subtyping approaches exist, with one emerging approach being the use of neuroimaging features (Hong et al., 2020).

From a clinical perspective, this heterogeneity is also mirrored in the pharmacological management of ASD. No single medication has been shown to improve the core social symptoms of ASD reliably, and most randomized controlled trials have failed to show superiority to placebo, primarily due to the extensive heterogeneity in ASD (Hellings, 2023). In fact, recent pharmacologic guidelines for ASD therapy recommend a symptom-targeted management strategy to address comorbid conditions (e.g., depression, anxiety, aggression) (Manter et al., 2025). This leads to the use of different medication classes from antipsychotics to antidepressants, resulting in complex polypharmacy treatments that may dramatically differ from patient to patient (Ritter et al., 2021). This clinical reality suggests that ASD reflects a family of overlapping neurodevelopmental trajectories that interact with multiple neuromodulatory systems. A key challenge for precision psychiatry in autism is therefore to identify stable, biologically grounded dimensions that can help explain why different individuals require different pharmacological strategies.

This clinical heterogeneity appears to be rooted in structural biology. Converging evidence from genetic studies also supports the notion of heterogeneity in autism, with a heterogeneous set of genes conferring risk (Rylaarsdam and Guemez-Gamboa, 2019). Many of these genes converge on neurodevelopmental pathways involved in neuronal growth and synaptic formation. Recent studies further link the polygenic risk score for autism to macro- and microscopic brain structure, associated with widespread cortical thickness and neurite density measures (Khundrakpam et al., 2020; Gu et al., 2025). Beyond polygenic scores, transcriptomic studies have shown that the spatial pattern of cortical thickness differences in autism closely follows the cortical expression of synaptic and downregulated ASD-related gene sets, directly linking genetic vulnerability to macroscopic anatomical organization (Romero-Garcia et al., 2019). Given that the genetic heterogeneity of autism appears to shape spatially organized differences in brain structure, neuroimaging offers a suitable tool for capturing the biologically grounded dimensions of heterogeneity.

A common strategy in neuroimaging-based subtyping has been the application of discrete clustering algorithms, which assumes that the neuropathology of a subgroup can be represented by a single, representative brain state (Duan et al., 2025). This premise, however, is inconsistent with the understanding of brain function, wherein multifaceted behaviors, such as those perturbed in ASD, arise from the dynamic interaction of large-scale, distributed neural systems (Vissers et al., 2012). The attempt to reduce this complex, system-level interplay to a unitary brain state for each individual is a significant oversimplification. This has motivated a shift towards dimensional models, which can better accommodate this biological complexity (Duan et al., 2025). Rather than assigning individuals to a single class, this approach quantifies the contribution of several underlying latent factors for each participant. This yields an individual-specific profile of factor loadings, which offers a more granular representation of neuropathology and establishes a more robust empirical basis for developing personalized intervention. One of the promising works in the dimensional approach is a work by Tang et al. (2020). The study identified three latent factors in autism derived from resting-state functional Magnetic Resonance Imaging (rsfMRI)-based Functional Connectivity (FC) using a Bayesian approach. Moreover, these factors were also found to correlate with different behavioral phenotypes (Tang et al., 2020). This dimensional, latent-factor framework, therefore, provides a realistic foundation for parsing ASD heterogeneity. This motivated its adoption in the present study to support more personalized characterization of ASD.

Notably, the implementation of FC is limited by its intrinsically dynamic nature; FC measures are known to be influenced by numerous factors, such as cognitive state, in-scanner motion, and other transient factors (Power et al., 2012; Noble et al., 2019). These factors may compromise the stability of the subtypes derived using FC. In contrast, brain morphometric features offer more stable measures for subtyping (Hedges et al., 2022). A recent meta-analysis also found that individuals with ASD displayed morphometric changes of the brain, such as decreased Gray Matter Volume (GMV) in the left cerebellum and increase GMV in the right precentral gyrus (Guo et al., 2024). The changes were also found in other morphometric features. Postema et al. (2019) found altered cortical thickness asymmetry in ASD especially in medial frontal, orbifrontal, cingulate, and inferior frontal regions. Moreover, Sha et al. (2022) showed that, compared with typical features of small-world architecture in controls, the ASD sample showed significantly altered average asymmetry of networks involving the fusiform, rostral middle frontal, and medial orbitofrontal cortex. These findings underscore the potential of morphometric features to characterize the subtypes in ASD.

Moreover, morphometric features derived from structural neuroimaging have also been employed for dimensional subtyping. For instance, Aglinskas et al. (2022) utilized anatomical brain volumes within a contrastive variational encoder framework to identify latent structural factors. Other work, such as Ecker et al. (2010), applied support vector machines to various area-based structural measures, including cortical thickness, concavity, and curvature. However, these approaches typically treat regional metrics independently. To create a structural analogue to the connectivity-based framework of Tang et al. (2020), a measure that quantifies inter-regional similarity based on a profile of structural properties is required.

Addressing this gap, a recent study proposed advanced methods to parameterize structural connectivity based on multiple structural features (Sebenius et al. 2023). The measure, called Morphometric Inverse Divergence (MIND), estimates similarity between cortical areas based on the divergence between their multivariate distributions of multiple morphometric features. This measure yielded more reliable morphometric similarity compared to prior approaches of morphometric similarity networks. Therefore, the use of MIND as a morphometric measure to explore the dimensional subtypes of autism is a promising approach to complement the findings from functional features.

The present study, therefore, explored the use of MIND as a morphometric measure in conjunction with the latent factors approach to explore the heterogeneity of ASD. We were also interested in investigating how the factors derived in ASD are different from the pattern observed in Healthy Controls (HC). We utilized a Bayesian approach as performed in the previous study (Tang, et al., 2020) to identify factors in individuals with ASD using the MIND measure from the participants. To provide a baseline for these factors, we compared these factors with the across-person average MIND from HC. Moreover, we also extended this analysis to FC measures, to also explore the similarity and mismatch of factors derived from MIND and FC in ASD participants. To further interpret the factors, we correlated all the factors with behavior measures. Given the nature of FC being influenced by numerous factors, and given the proven relationship between structural brain measures and ASD, we hypothesized that factors derived from morphometric measures (i.e., MIND) would provide a more behaviorally relevant account of ASD heterogeneity compared to those from functional measures (i.e., FC). Finally, we aimed to explain the biological underpinning of those factors. To this end, we map these factors into previously defined neurotransmitter receptor atlases (Hansen et al. 2023). Finally, we discussed our findings on morphometric and functional factors derived to dimensionally subtype ASD and how they could contribute toward the development of targeted therapies and biomarkers.

## 2. Methods

The main aim of the current study was to explore the autism spectrum disorder in terms of their morphometric features using MIND measure based on a dimensional approach. Firstly, both structural and functional MRI of individuals with autism from the Autism Brain Imaging Data Exchange II (ABIDE-II) dataset was acquired. The ABIDE-II is a dataset including 1114 individuals collected from 19 different sites. The dataset includes a total of 521 individuals with ASD and 593 control individuals (Di Martino et al., 2014; 2017). We then proceeded with preprocessing the structural and functional data for each participant. Next, we computed the MIND matrix of each participant representing their morphometric features. This matrix informs the similarity of the morphometric features between brain areas. Similarly, we also calculated the FC matrix of each participant from the resting-state functional MRI data, representing the similarity of activation patterns between brain areas. Finally, we implemented a Bayesian approach used by Tang et al. (2020) on both MIND and FC obtaining three different factors each for each modality. Note that here, unlike Tang et al. (2020) who derived the factors of ASD from FC normalized by HC, we only used MIND and FC from ASD without normalization. Notably, this Bayesian framework also provides loading of each participant on each of these factors depending on how similar the factor pattern is to their functional connectivity and structural similarity patterns. The loadings were subsequently used to calculate the correlation between factors and behavior. Finally, to expand on our findings, we reduced each of the factors from both MIND and FC into a single principal component for each factor and regressed it onto 19 different neurotransmitter density maps.

To further validate the factors derived solely from ASD participants, we compared these factors with group-average MIND and FC from the HC in ABIDE-II dataset. We also included HC from the Human Connectome Project (HCP) to validate and compare findings across the healthy controls in the two datasets. We detailed each of the methods below.

### 2.1 Participants

Similar to Tang et al. (2020), both structural and resting-state functional MRI brain scans were obtained from the ABIDE-II dataset (Di Martino et al., 2014; 2017). Our study involved 835 participants from ABIDE-II in total. Among them, 368 were ASD participants (*N_males_* = 255, *M_age_* = 15.34, *SD_age_* = 9.10) and 467 were control group individuals (*N_males_* = 379, *M_age_* = 15.03, *SD_age_* = 8.76). The data was collected from multiple sites, each site had a slightly different inclusion criteria; however, in general, most sites were able to confirm the diagnosis using the Autism Diagnostic Observation Schedule or the Autism Diagnostic Interview Revised. For more details about the dataset, please refer to their official website: https://fcon_1000.projects.nitrc.org/indi/abide/abide_II.html.

### 2.2 Analysis

#### 2.2.1 Functional MRI Data Preprocessing and FC Computation

For the preprocessing of the rsfMRI data, we followed the CBIG pipeline for fMRI preprocessing through the following steps proposed by Thomas Yeo’s lab which can be found in the following GitHub repository: (https://github.com/ThomasYeoLab/CBIG/tree/master/stable_projects/preprocessing/ CBIG_fMRI_Preproc2016). First, we removed the first four frames of the data for each participant. We then performed slice timing correction using FSL 6.0 (https://fsl.fmrib.ox.ac.uk/fsl/docs/). We used the slice order provided by the labs that collected each of the sub-datasets in the ABIDE-II dataset or, in case the slice order was not specified, we used the default slice order of the machines. Motion correction using rigid body translation and rotation was also applied using FSL. We then aligned the functional data to the structural images using boundary-based registration in `FsFast` from FreeSurfer. The command of `fsl_motion_outliers` was then used to calculate Framewise Displacement (FD) and Differentiated Signal Variance (DVARS). Volumes with FD bigger than 0.2 mm or DVARS bigger than 50 were excluded as outliers. Linear regression of multiple nuisance variables was then applied. The nuisance regressors consisted of (a) a vector of linear trend; (b) six motion correction parameters; (c) averaged white matter signal; (d) averaged ventricular signal; (e) global signal; (f) temporal derivatives of (b)-(e). The data were subsequently band-pass filtered (0.009 Hz ≤ *f* ≤ 0.08 Hz), projected onto the FreeSurfer fsaverage6 surface, smoothed with a 6mm full-width half maximum (FWHM) kernel, and down-sampled to the fsaverage5 surface space. Finally, we parcellated the cleaned functional data according to Schaefer400 parcellation (Schaefer et al., 2018). The FC was calculated as a Pearson correlation between pairs of regions, resulting in a 400×400 matrix for each individual. The final sample after the preprocessing and exclusion was 368 participants in the autism group and 467 participants in the control group.

#### 2.2.2 Structural MRI Data Preprocessing and MIND Computation

Participants’ T1-weighted MRI brain scans were reconstructed using the `recon-all` command in Freesurfer 7.4. We then segmented the brain scans of each participant surface using the Schaefer400 brain atlas (Schaefer et al., 2018). Cortical thickness, volume, area, curvature and sulcus depth were calculated for each of the vertices in the brain. These morphometric features were then used to calculate MIND measure for each pair of regions (Sebenius et al., 2023). Briefly, this measure treats each region in the brain as a distribution of the vertices. The Kullbach-Leibler divergence was then computed for each pair of regions. We then z-normalized these values, and the resulting matrix is a 400×400 matrix, similar to that of the FC matrix.

#### 2.2.3 ASD Factor Identification

To identify the factors, we fully adopted the Bayesian framework by Tang et al. (2020) and Yeo et al. (2015). To increase the utility of the code, we transformed it from a C code into a python code. We checked the outputs from the C and Python codes using the same datasets to validate that the Python code works as intended. We found that the outputs between them are highly similar (see Figure 6).

The Bayesian framework that was applied to identify the factors was based on the Latent Dirichlet Allocation (LDA) model, introduced by Blei et al. (2003). It is an unsupervised generative machine learning model designed for text processing. The main goal is to identify the distribution of topics within a collection of documents by analyzing the frequency of words in each document and identifying patterns of word co-occurrence across the entire corpus. The framework was then also extended to discover latent factors in Alzheimer’s Disease (Zhang et al., 2016) and cognitive tasks (Yeo et al., 2015).

In the present study, the approach was implemented on both FC and MIND matrices of ASD individuals to identify functional and morphometric factors, respectively. In brief, the method works by analyzing the connectivity between pairs of brain regions across participants and clustering them into distinct factors based on their commonality. Moreover, each individual is assigned a factor loading to indicate how closely their FC and MIND are aligned with the functional and morphometric factors, respectively. These loadings represent the dimensional clustering of the ASD; instead of assigning individuals to a single class, this approach quantifies the contribution of several underlying latent factors for each participant. The individual loadings are also inline with the research trend to develop personalized diagnosis and treatment.

To interpret the factors, we adopted the terms used by previous study (Tang et al., 2020), which are hyper-connectivity for positive connections and hypo-connectivity for negative connections in the functional factors derived from FC. For the morphometric factors, we used hypo- and hyper-similarity for negative and positive morphometric similarity derived using MIND. For the technical details on how the method was implemented, please refer to the original study by Tang et al. (2020).

#### 2.2.4 Comparison with Healthy Control

Notably, we compared the functional and morphometric factors identified from the ASD individuals with factors derived from Healthy Control (HC). To identify the functional and morphometric factors from the HC, we simply averaged the FC and MIND of the healthy individuals, respectively. To compare the factors between ASD and HC, we took the lower triangle of the factor matrices and computed the Pearson correlation between two factors. For example, to compare the functional Factor 1 from ASD individuals with the factor from the HC, we computed the Pearson correlation between the vectorized lower triangle of functional Factor 1 matrix from ASD and the vectorized lower triangle of factor from the HC.

For validation, aside from the healthy individuals in ABIDE-II dataset, we also employed data from the Human Connectome Project Young Adult (HCP-YA) dataset from 100 individuals (*N_males_* = 46, age *M_age_* = 29.1, *SD_age_* = 3.7 years) as a controlled test of a separate sample HC (Van Essen et al., 2012). We downloaded the minimally preprocessed structural MRI data from the HCP-YA dataset. The cortical grey matter was then parcellated to the Schaefer 400-parcel atlas to ensure consistency with the current study, before computing the MIND matrices. For rsfMRI, we used the minimally preprocessed, ICA-FIX–denoised functional data provided by the HCP (Glasser et al., 2013; Salimi-Khorshidi et al., 2014). For each of the four rsfMRI runs per participant, time series were extracted using the Schaefer 400 atlas, pairwise FC matrices were computed, Fisher r-to-z transformed, and subsequently averaged across runs.

#### 2.2.5 Association of Factors with Behavioral Measures

To go beyond identifying factors in ASD individuals and presenting the factor loadings, we investigated their associations with behavioral measures and neurotransmitter densities.

For the first association, we extracted the factor loadings of ASD individuals to each of the factors. This gave three loadings for each participant (one for each factor). We then computed the Pearson correlation of this loading with different behavioral measures. This analysis was repeated in all morphometric and functional factors. Importantly, we used the same set of behavioral measures as used in the previous study, Tang et al. (2020). This set consisted of the Social Communication Questionnaire (SCQ) and the Repetitive Behaviors Scale – Revised (RBSR). The SCQ is a widely used measure for screening for autism. Research in general has reported considerable validity and reliability for the test (Hollocks et al., 2019). Developed by Rutter et al. (2003), the test consists of three different sub-factors: reciprocal social interaction, communication, and restricted. The RBSR, on the other hand, is a scale developed to test individuals on their repetitive and restrictive behaviors (Bodfish et al., 2000). The test consists of five different sub-factors: stereotypic behaviors, self-injury, compulsive, ritualistic, and repetitive behaviors. Similar to the SCQ, the RBSR scale was reported to have quite good validity and reliability (McDermott et al., 2020). We also used the social responsiveness scale (SRS; Brooker & Starling, 2012). The test was developed for the diagnosis and assessment of autism spectrum disorder in children and has demonstrated a strong sensitivity score (91%; Aldridge et al., 2012).

#### 2.2.6 Association of Factors with Neurotransmitter Densities

For the association with the neurotransmitter densities, we used the neurotransmitter density distributions of 19 different neurotransmitters extrapolated from Positron Emission Tomography (PET) data of 1200 healthy individuals and published by Hansen et al. (2023). The neurotransmitter map is available for each brain region in Schaefer400 atlas, the parcellation we used in this study. We then investigated the association between the neurotransmitter maps and the factors in order to identify the corresponding neurotransmitters of each factor. Because each latent factor is represented as a symmetric connectivity matrix while the neurotransmitter maps are region-wise vectors, we reduced each factor to a single summary vector using Principal Component Analysis (PCA). PCA extracts the dominant pattern of connectivity variation across regions, resulting in a vector that summarizes the connectivity patterns of each brain area. This vector can then be directly compared with the neurotransmitter distributions. We then regressed the 19 neurotransmitter distribution maps on each of the vector summaries from functional and morphometric factors derived in ASD individuals. We then took the regression weights of each neurotransmitter and interpreted them as the contribution of different neurotransmitters to each factor. Moreover, after running a PCA on the average HC FC and MIND data we also ran this regression analysis on the resulting PC to compare the underlying neurotransmitters between factors derived from ASD individuals and HC. In this case, the PCA was applied on the functional and morphometric factors from the healthy individuals (group-average FC and MIND, respectively). Notably, we used HC datasets from ABIDE-II and HCP-YA.

Because brain data exhibits high spatial autocorrelation, for the association between the factors and the neurotransmitters, we decided to run a spin permutation hypothesis test, rather than depending on theoretical p-values to test our hypotheses. This method has been argued to be much more realistic when considering brain data and avoids over estimation of the significance levels (Alexander-Bloch et al., 2018).

#### 2.2.7 Data and Code Availability

The ABIDE-II data used in this analysis are available on the International Neuroimaging Data-Sharing Initiative website: https://fcon_1000.projects.nitrc.org/indi/abide/abide_II.html. The HCP-YA data can be found at the following link: https://balsa.wustl.edu/project?project=HCP_YA. The analysis scripts are available on the following GitHub repository: https://github.com/ajn3333/Latent_Factor_ASD/tree/main.

## 3. Results

We organized the results into three different sections. We first presented the neural profiles of the identified functional and morphometric factors. Second, we showed how they differ between ASD and HC. Third, we presented how these factors were correlated with behavioral measures. Finally, we identified the biological underpinnings of these factors in terms of their neurotransmitter correlates.

### 3.1 The Neural Profiles of Functional and Morphometric Latent Factors and the Individual Loadings

The functional and morphometric factors derived from ASD individuals are depicted in Figure 1. For the functional factors (Figure 1 A), we found clear patterns of hypo- and hyper-connectivity in different brain regions. Generally, we found that these factors were similar to the factors identified by the previous study (Tang et al., 2020; see also Figure 6). The first factor (the left figure) involved hypo-connectivity within the default mode, visual, and somatomotor areas, as well as hyper-connectivity between somatomotor and visual areas, and between visual and default mode areas. Moreover, the second factor (the middle figure) was characterized by the hyper-connectivity within the somatomotor, visual, dorsal attention, and default mode areas. Interestingly, this neural profile is reversed in the third factor (the right figure), where it showed hypo-connectivity within the somatomotor network and between somatomotor areas and visual areas. For the morphometric factors (Figure 1B), we found hypo-similarity within visual and somatomotor networks, as well as between these networks and all other networks in the first factor (left factor). Conversely, the second and the third factors showed more hyper-similarity between different brain networks, especially involving visual and somatomotor networks.

**Figure 1.**
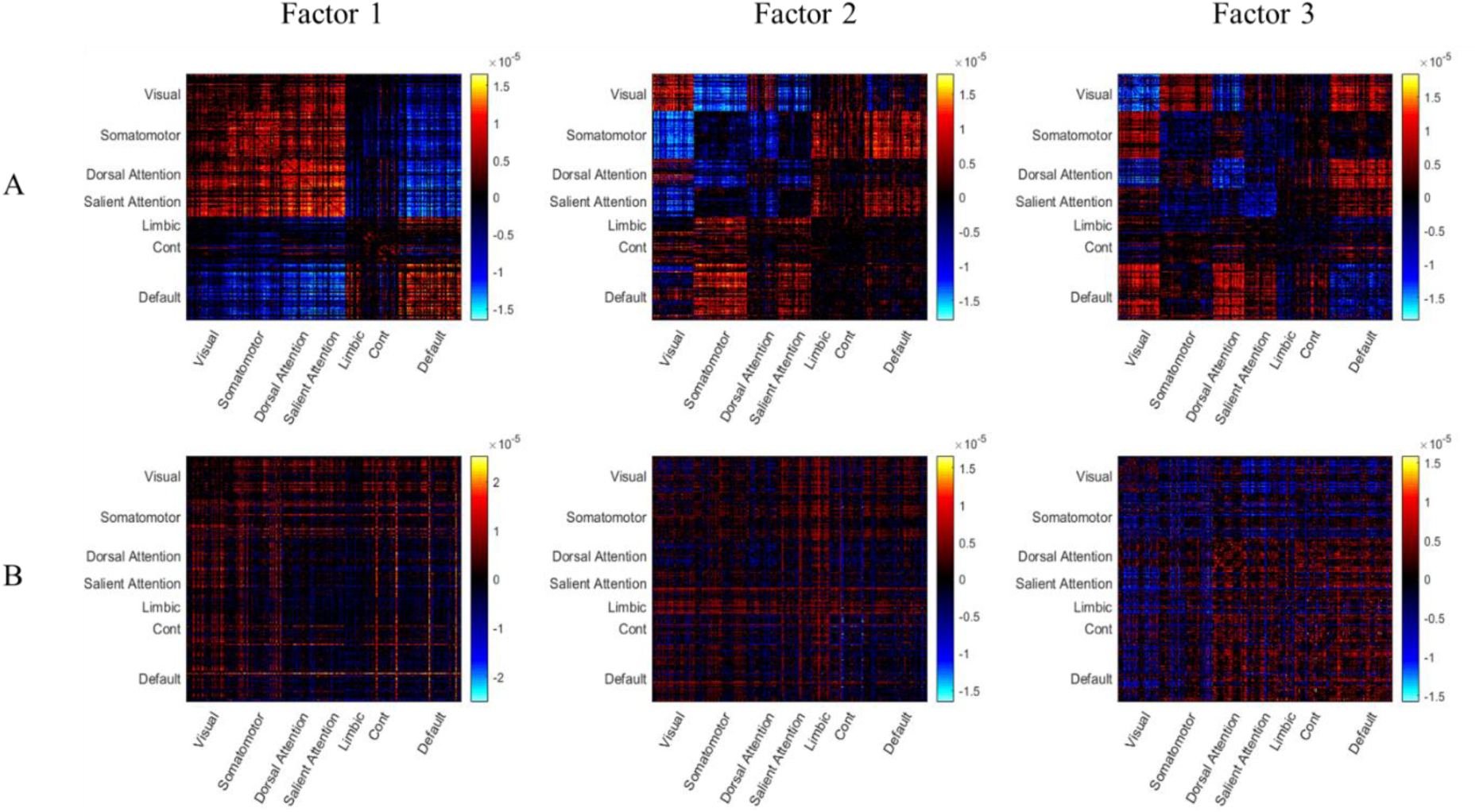
Latent factors of functional connectivity and morphometric similarity in ASD. The three latent factors derived from (A) Functional Connectivity (FC) and (B) Morphometric Inverse Divergence (MIND) data from ASD participants. Matrices represent the factor patterns, with brain regions grouped by large-scale functional networks. Red and blue colors indicate patterns of hyper- and hypo-connectivity (for FC) or hyper- and hypo-similarity (for MIND), respectively.

To visualize the individual-level distribution of these factors, we plotted the participant loadings for both modalities on Figure 2. For the functional factors (Figure 2A), the loadings from the ASD participants were widely distributed across the entire space. This spread indicates a high degree of inter-individual heterogeneity, consistent with the findings of Tang et al. (2020), where different individuals show strong preferences for each of the three factors, as well as many individuals exhibiting hybrid profiles (i.e., dots in the center of the triangle). In contrast, the morphometric factor loadings (Figure 2B) displayed a more constrained pattern. The distribution was not uniform; instead, the loadings were heavily concentrated along the simplex edge connecting Factor 1 and Factor 3. This pattern implies that the structural variation is dominated by a trade-off between MIND factor 1 and MIND factor 3, with MIND factor 2 representing less contribution to individual pathology.

**Figure 2.**
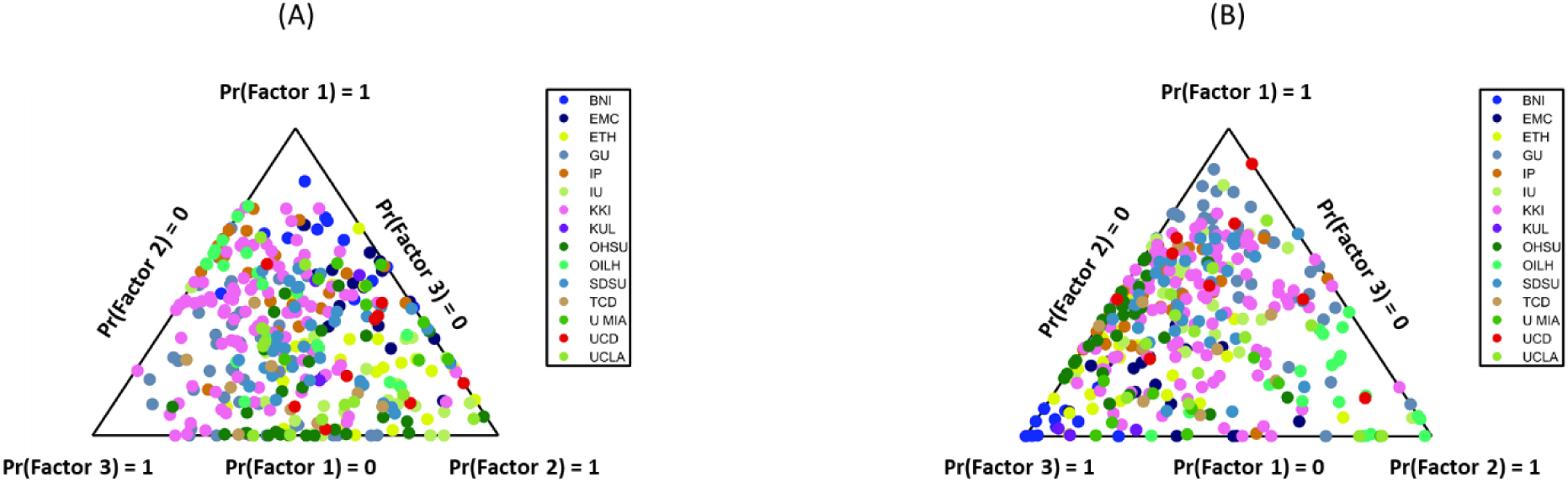
Individual factor loadings for functional and morphometric factors in ASD. Simplex plots illustrate the distribution of individual ASD participant loadings for (A) the three functional factors (FCF1-3) and (B) the three morphometric factors (MINDF1-3). The wide spread in (A) indicates high functional heterogeneity, while the constrained pattern in (B), concentrated along the F1-F3 edge, shows a more specific structural profile with a reduced contribution from Factor 2. Each dot represents one participant, colored by the acquisition site.

### 3.2 ASD Showed Weaker Correspondence Between Functional and Morphometric Factors Compared to HC

It is important to note that unlike ASD factors which were derived using Bayesian approach, the factors for HC were simply the average of FC or MIND matrices across individuals. We then compared these factors by correlating the lower triangle of the corresponding matrices. Generally, the results revealed a significant correlation between the ASD factors and HC group-average connectivity (see Figure 3). Specifically, functional factor 1 (*r =* -.36, *p* < .001) and factor 3 (*r* = .48, *p* < .001) revealed a moderate correlation with the HC group-average FC, while factor 2 showed weak correlation (*r* = -.01, *p* < .001). The MIND factors exhibited a negative weak correlation between the factor 2 and the HC group-average MIND (*r* = .06, *p* < .001), a moderate negative correlation for factor 3 (*r* = -.24, *p* < .001), and a moderate positive correlation for factor 1 (*r* = .27, *p* < .001).

**Figure 3.**
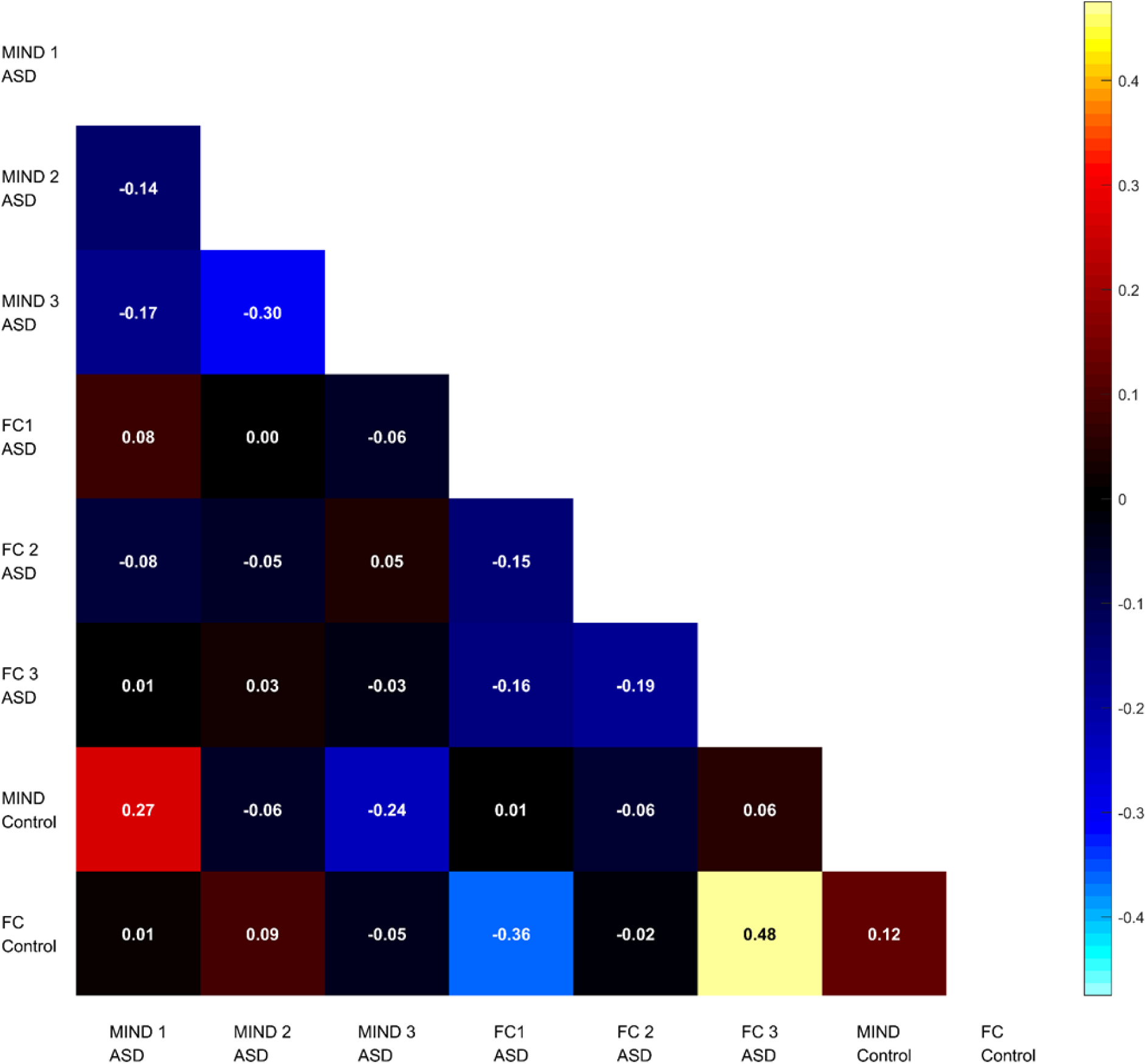
Correspondence between functional and morphometric factors in ASD and HC. Pearson correlation matrix illustrating the associations between the three functional (FC) and three morphometric (MIND) factors in ASD, as well as the group-average FC and MIND patterns from HC. The matrix highlights the weak negative association between structural and functional factors within the ASD group, which contrasts with the positive structure-function correspondence observed in the HC group.

Moving to the functional and morphometric factors correspondence, we found that ASD individuals showed weak negative to no association between functional and morphometric factors. Interestingly, the association tended to be positive (*r* = .12, *p* < .001) in HC (see Figure 3). This finding contributes to the understanding of the differences in terms of structure and function coupling in ASD and HC, showing that in ASD, the coupling between them was weaker compared to HC.

### 3.3 The Loadings of Morphometric Factors Were More Associated with Behavioral Measures Compared to the Functional Factors

To interpret the factors in terms of their behavioral correlate, we correlated the individual factor loadings to different behavioral scores related to ASD. On one hand, we found no significant correlation between the functional factor loadings of the individuals and their behavioral scores. On the other hand, the morphometric factor loadings extrapolated from the MIND showed significant correlation with the behavioral scores. The SCQ scale was correlated with the loadings of factor two (*r* = .29, *p* < .005) and factor three (*r* = -.31, *p* < .005). A significant correlation was found also between the SRS scale and the loadings of factor one (*r* = -.16, *p* < .005) and factor two (*r* = .16, *p* < .005). No significant correlations were found between the RBRS scores and loadings from any factors.

### 3.4. The Neurotransmitter Correspondence of the Factors

For the last analysis, we performed a regression model to find the neurotransmitter correspondence of different factors from ASD and of HC group-average connectivity. The results were depicted in Figure 4. For the functional factors, we found that the norepinephrine transporter (NET) receptor distribution seems to be significantly positively associated with the ASD factors, but not with the HC. For the morphometric factors, we found a significant positive relationship between the CB1 receptor distribution with the healthy factors, but not with the ASD factors. These findings indicate that NET and CB1 receptors might be related to ASD, in terms of their neural functions and structures, respectively. The roles of NET and CB1 receptors in healthy factors were also tested with the HCP-YA dataset. The result aligned well with the HC analysis from the ABIDE II dataset, where CB1 receptor was associated with the group-average MIND. Interestingly, for the functional principal components, the result suggested a negative relationship between HC group-average FC and the NET receptor distribution, contrasting the finding from the HC in ABIDE II datasets. Notably, here we emphasize that the two datasets differ in terms of their age distribution.

**Figure 4.**
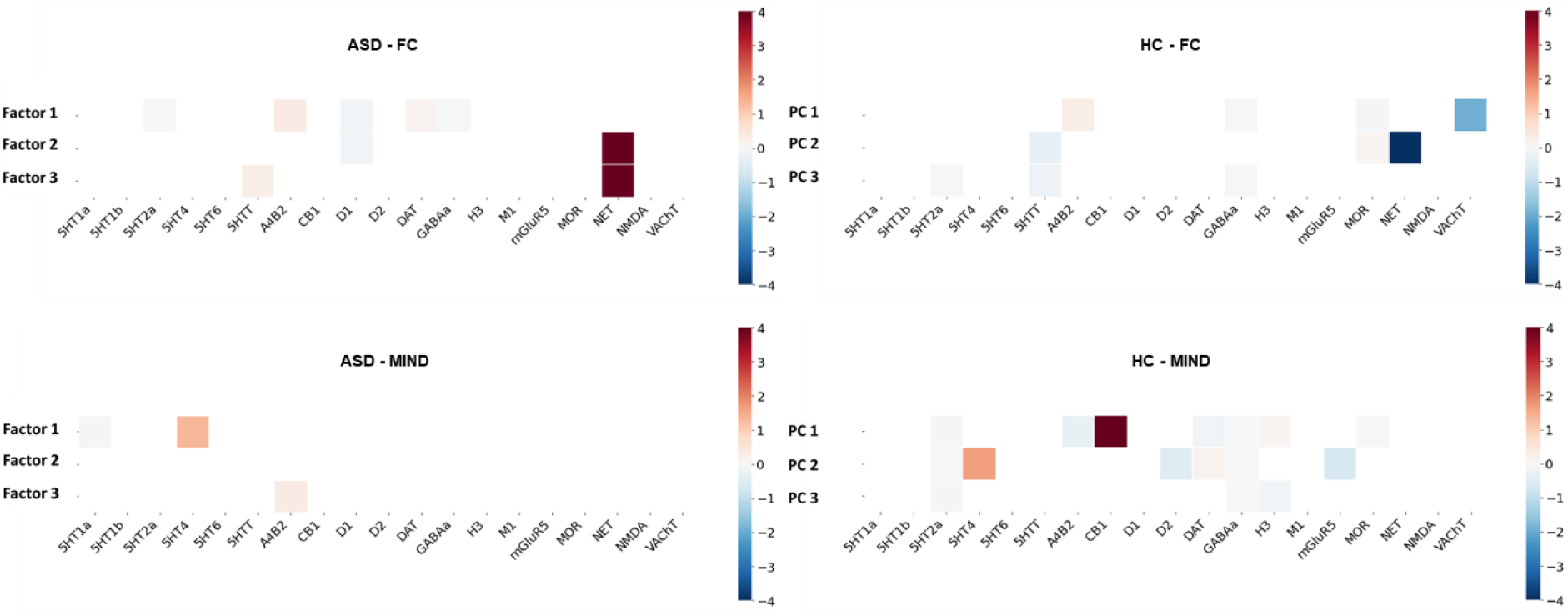
Neurotransmitter receptor profiles of functional and morphometric factors in ASD and HC. Heatmaps showing the regression beta coefficients from associating 19 neurotransmitter receptor density maps with the functional (top row) and morphometric (MIND, bottom row) factors. Results are shown for both the ASD factors (left column) and the HC group-average patterns (right column). The results highlight distinct neurochemical signatures, particularly the strong association of the norepinephrine transporter (NET) with ASD functional factors and the association of the CB1 receptor with the HC morphometric pattern.

**Figure 5.**
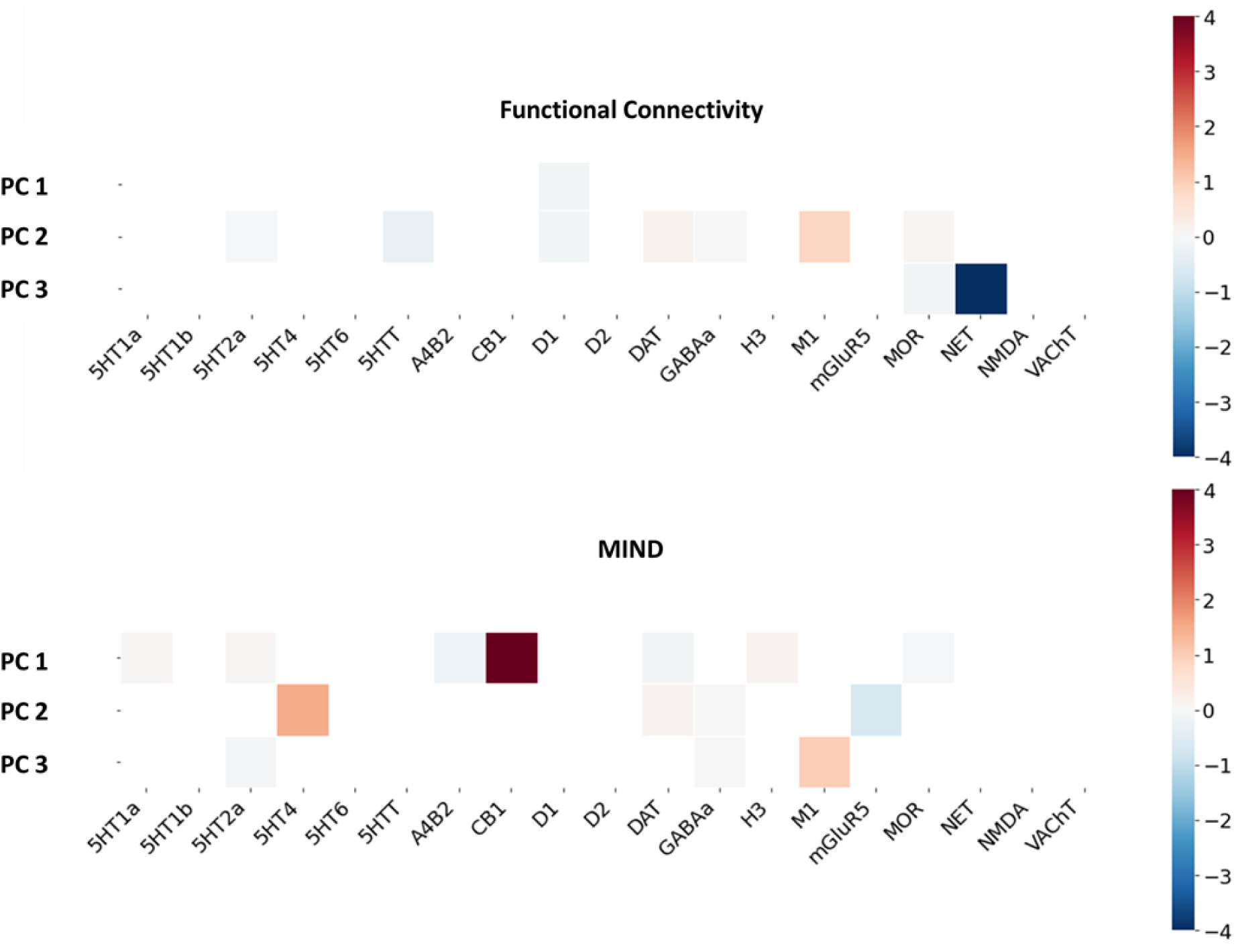
Validation of neurotransmitter associations in the HCP-YA dataset. Heatmaps showing regression beta coefficients associating neurotransmitter receptor maps with the principal components of group-average Functional Connectivity (top) and MIND (bottom) from the independent HCP-YA validation cohort. These results serve to validate the findings from the ABIDE-II HC group, confirming the association of the CB1 receptor with the healthy morphometric profile.

**Figure 6.**
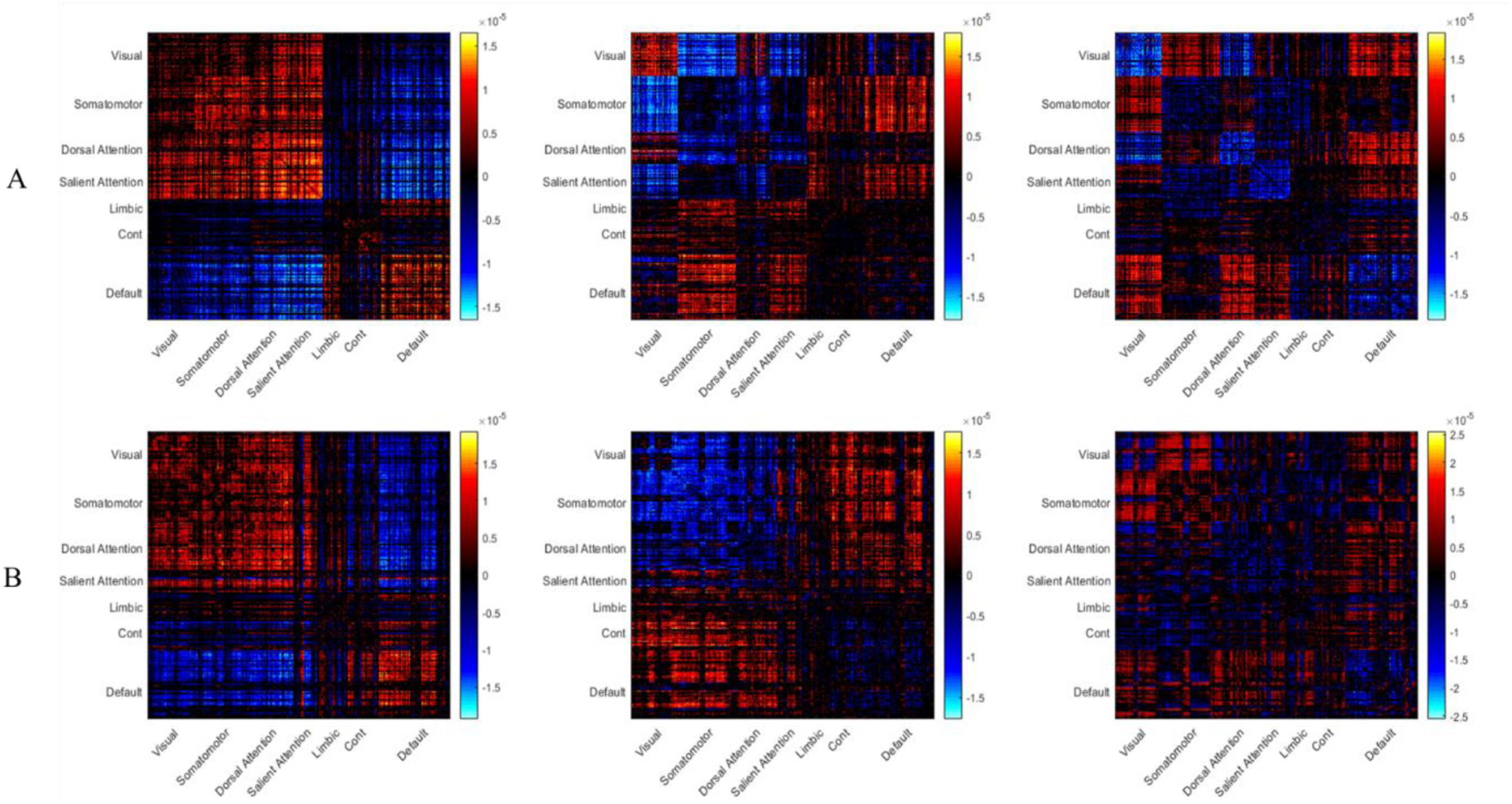
Comparison of FC factors derived with Factors derived by Tang et al. (2020). A visual comparison between the three functional connectivity (FC) factors derived in the present study (A) and the original three FC factors reported by Tang et al. (2020) (B). The qualitative similarity between the factor patterns validates the successful replication of the original LDA methodology and the transformation from C- to python-based scripts.

## 4. Discussion

In this study, we applied a dimensional framework to characterize the heterogeneity of autism using the functional connectivity and morphometric similarity networks. Although FC and MIND factors were moderately correlated, each modality yielded distinct dimensional subtypes in autism, underscoring the separable functional and structural correlates of the variation. Notably, unlike FC, only MIND factors were highly correlated with social and anxiety symptoms of ASD, suggesting that morphometric similarity networks may capture more trait-like features of autism. The MIND factor profiles were comparable across the healthy controls of two cohorts (ABIDE-II and HCP-YA), indicating the stability of this approach. Finally, neurotransmitter mapping revealed neuromodulatory correlates for these dimensions: NET linked to FC-based factors, and CB1 receptor distribution linked to MIND-based factors, providing a biological and a possible clinical interpretation of the dissociation between functional and structural dimensional subtyping.

First, we found that factors derived from FC were highly similar with the ones identified by Tang et al. (2020), emphasizing the reproducibility of the research findings. The analysis using MIND data revealed a global pattern difference between functional and morphometric factors. The difference across functional factors was more about network-specific connectivity, aligning with the established ASD literature that highlights atypical coupling in systems such as the default mode and salience networks (Abbott et al., 2016). These factors represent the complex, downstream functional expression of the disorder represented by the network-wise connectivity.

Differently, the morphometrical factors were differentiated by their global composition of hypo- and hyper-similarity. Factor 1 was dominated by hyper-similarity, factor 2 was a mix, and factor 3 tended toward hypo-similarity. This suggests the MIND-derived factors may be capturing a more fundamental and trait-like biological profile, which is inherently more stable than the state-dependent nature of functional connectivity (Betzel & Bassett, 2017). In addition, this interpretation aligns with genetic studies in ASD, which indicate that genes involved in ASD converge on neurodevelopmental pathways (Rylaarsdam and Guemez-Gamboa, 2019) that subsequently manifest as a spatially specific pattern of cortical thickness differences (Romero-Garcia et al., 2019). Moreover, one of the theories of the root cause of ASD posits that an early and rapid brain overgrowth (Courchesne et al., 2003) may be driven by widespread deficits in synaptic pruning, which is the normal developmental clean-up of unnecessary neural connections (Tang et al., 2014; Eltokhi et al., 2020). Such a process would result in a cortex that is less neuroanatomically specialized and more morphometrically undifferentiated. The widespread hyper-similarity observed in MIND factor 1 could therefore be a large-scale, neuroanatomical signature of this proposed etiological mechanism. Such a singular, global structural bias could then be the ‘upstream’ driver that gives rise to the multiple, complex, and network-specific functional disruptions observed in the FC factors.

Moreover, we also found that the loadings from the morphometric factors were better associated with ASD-related behavior compared to functional factors. This finding provides further evidence for our interpretation of the structural factors as upstream biological traits. Core ASD behavioral phenotypes are stable, long-term traits, and to effectively predict them, the underlying brain measure must also be a reliable, trait-like marker (Lombardo et al., 2019). Functional connectivity, while powerful, is intrinsically dynamic and has well-documented challenges with test-retest reliability, making it susceptible to transient, non-pathological ‘noise’ (Noble et al., 2019). This inherent variability may obscure stable brain-behavior relationships. In contrast, morphometric measures are highly reliable (Hedges et al., 2022). This connects directly to our prior point: the MIND factors, by capturing the global proportion of structural similarity, are indexing this stable and foundational organization. Our results suggest it is this fundamental structural trait, not its noisy functional consequence, that serves as the more robust predictor of behavioral phenotypes.

Second, moving to the functional and morphometric factors correspondence, we found a positive association in the HC group (note that here for HC the factors were identified simply by taking the average FC or MIND matrices for all the participants). This is consistent with the fundamental neuroscience principle of structure-function coupling, where the brain’s structural architecture provides a foundation that constrains and shapes its functional dynamics (Park & Friston, 2013). In contrast, we found that ASD individuals showed weaker and mainly negative association between the MIND and the FC factors. This suggests a structure-function decoupling in the autistic brain, a finding that aligns with other recent reports of reduced structure-function coupling in ASD (e.g., Qing et al., 2024; Qiao, et al., 2025; Valenti et al., 2020). This decoupling may be a direct macro-scale consequence of the etiological theories discussed previously; if the structural foundation is less specialized due to widespread deficits in synaptic pruning (Tang et al., 2014), it may be incapable of effectively guiding its functional expression. The resulting functional patterns would then be less constrained by the underlying anatomy, leading to the exact decoupling we observed in our data.

Third, our analysis of the neurochemical correlates for these factors revealed a dissociation which may explain why the structural factors, and not the functional factors, were predictive of behavior. For the functional factors, we found a significant association with the NET in the ASD group, a relationship not observed in the HC group. This finding may seem contradictory with the other finding on behavior correlate: the functional factors themselves did not correlate with stable behavioral traits, yet they are tied to a neuromodulatory system (the noradrenergic system) that is widely implicated in ASD-related behaviors (Beversdorf, 2020). This apparent mismatch can be resolved by considering the state-like nature of both measures. The noradrenergic system governs moment-to-moment processes such as arousal, vigilance, and sensory-driven attention (Zhao, et al., 2022). Functional connectivity is itself intrinsically dynamic (Noble et al., 2019). It is therefore logical that the state-like functional factors of ASD are spatially tethered to a state-like neuromodulatory system. This may also explain why these factors failed to predict behavior; a highly variable arousal-dependent measure is an unsuitable proxy for the stable trait-like behavioral phenotypes of ASD, such as those measured by the SCQ and SRS.

In contrast, the morphometric factors, which did predict behavior, were linked to a trait-like biological signature. We found that the primary morphometric pattern in the HC group was significantly associated with the CB1 receptor, but this association was notably absent in the ASD factors. This is an important finding, as the endocannabinoid system (via CB1) is a fundamental regulator of neurodevelopment, primarily by guiding neural migration and initiating synaptic pruning (Harkany et al., 2007; Costello & Roche, 2025). The absence of this normal CB1-structure relationship in the ASD group provides a specific, neurochemical mechanism for the deficient pruning (Tang et al., 2014) and early brain overgrowth (Courchesne et al., 2003). This completes our central narrative: a failure in the CB1-mediated trait-building process leads to a stable atypical structural organization (our MIND factors), which in turn predicts the stable, long-term behavioral traits of ASD.

Furthermore, our validation analysis in the HCP-YA cohort underscores the stability of the morphometric-based findings. We found that the significant association between the CB1 receptor and the group-average MIND profile was replicated in both the ABIDE-II and the independent HCP-YA healthy control groups. In contrast, the neurochemical signature of functional connectivity proved inconsistent; the NET-FC relationship differed between the two HC cohorts. This functional instability is not surprising, given that FC is an intrinsically dynamic measure sensitive to developmental change. As we noted, the two cohorts differ in their age distribution, which likely accounts for this discrepancy. This divergence further reinforces our premise: for identifying stable, trait-like biological markers, stable structural measures such as MIND offer a more reliable foundation than their dynamic functional counterparts.

Another implication of these findings relates to the longstanding issues in pharmacological treatment of ASD. Clinical trials targeting various neuromodulatory systems have generally failed to achieve satisfying results in treating the core symptoms of ASD (Hellings, 2023), while they were successful in treating the co-occurring symptoms. An example of these medications is Atomoxetine, a NET inhibitor, which has been shown to improve attention- and arousal-related symptoms in these patients (Harfterkamp et al., 2012). Our results provide a potential mechanistic context for this pattern: the functional factors in ASD were related to NET, consistent with the high prevalence of attention-deficit/hyperactivity in ASD (Taurines et al., 2012) and with evidence that neuradrenergic modulation by targeting NET specifically affects these symptoms in ASD (Harfterkamp et al., 2012). In contrast, morphometric latent factors that predicted the core symptoms were related to CB1 receptor distribution, possibly reflecting the neurodevelopmental role of this receptor (Harkany et al., 2007; Costello & Roche, 2025). However, in a recent proof-of-concept randomized trial to evaluate the tolerability of cannabinoid treatment, no significant effect in treating core ASD symptoms were observed (Aran et al., 2021). While our findings do not imply any immediate clinical applicability, they raise the possibility that stable, biologically grounded latent factors, particularly those derived from morphometric similarity, may eventually help define subgroups with differential engagement of neuromodulatory systems. Such stratification could, in principle, guide more precise testing of interventions in future clinical trials, although substantial longitudinal and mechanistic validation will be required before any medication decisions can be informed by these dimensions.

This study is not without limitations. First, the neurotransmitter profiles were derived from ex-vivo and in-vivo PET atlases of healthy adults (Hansen et al., 2023). This indirect mapping is a significant assumption, and future studies using ASD-specific, in-vivo imaging are needed to confirm these specific neurochemical associations. Second, our analysis was restricted to the cerebral cortex, thereby omitting subcortical regions such as the striatum, thalamus, and amygdala, which are critically implicated in ASD neuropathology. Finally, our analysis only included a cross-sectional look at ASD. However, longitudinal studies are required to confirm their stability as genuine biological traits.

Future work should aim to address these points by applying this MIND-based framework to longitudinal, whole-brain data. The translational potential of this approach is that these stable and behaviorally relevant factors could serve as a method for stratifying individuals in clinical trials to predict who may respond to specific interventions.

In conclusion, this study provides a direct comparison of functional and morphometric latent factors in ASD, revealing a compelling dissociation between the two. We demonstrate that stable factors of brain structure, derived using the MIND methodology, are significantly linked to the core behavioral traits of autism – a connection not observed for their functional counterparts. The observed structure-function decoupling in the ASD group, combined with the identification of a plausible neurodevelopmental (CB1) pathway for the structural factors, points to a fundamental alteration in the brain’s organizational principles. These results advocate for the use of advanced and reliable morphometric measures as a powerful and necessary path toward developing the stable and biologically grounded biomarkers required for personalized medicine in autism.

## Author Contribution

AAl: data curation, methodology, software, formal analysis, visualization, writing – original draft, AAb: supervision, data curation, methodology, software, formal analysis, writing – review and editing, DK: supervision, methodology, software, formal analysis, writing – original draft, writing – review and editing

## Declaration of Competing Interest

None

## Acknowledgment

DK was supported by the Young Researchers’ Fellowship from the University of Oldenburg and by a grant from the German Research Foundation (DFG) to Andrea Hildebrandt (HI 1780/7-1) and Carsten Gießing (GI 682/5-1) as part of the DFG priority program “META-REP: A Metascientific Programme to Analyse and Optimise Replicability in the Behavioural, Social, and Cognitive Sciences” (SPP 2317). AAb was funded by DFG as part of the Research Training Group (RTG)-2783 (Project ID: 456732630).

Data were provided in part by the Autism Brain Imaging Data Exchange (ABIDE) II. The primary funding source for ABIDE II was the National Institute of Mental Health (NIMH) grant 5R21MH107045.

Data were provided in part by the Human Connectome Project, WU-Minn Consortium (Principal Investigators: David Van Essen and Kamil Ugurbil; 1U54MH091657) funded by the 16 NIH Institutes and Centers that support the NIH Blueprint for Neuroscience Research; and by the McDonnell Center for Systems Neuroscience at Washington University.

## Notes

https://fcon_1000.projects.nitrc.org/indi/abide/abide_II.html

https://github.com/ajn3333/Latent_Factor_ASD/tree/main

### Competing Interest Statement

The authors have declared no competing interest.

